# Custom-Built Electrodes Perform Comparably to a Discontinued Commercial Electrode for Neuromuscular Electrical Stimulation in Mice

**DOI:** 10.1101/2025.04.30.651500

**Authors:** Bana H. Odeh, Amanda L. Wellman, Michael Ameye, Zachary Atwood, Luke Gray, Aiswarya Saravanan, Havish Poluru, Morium Begam, Takako I. Jones, Renuka Roche, Joseph A. Roche

## Abstract

**Objective:** A commercial electrode used for transcutaneous neuromuscular electrical stimulation (NMES) in mice is no longer available. In response, we developed two low-cost, customizable alternatives—a 3D-Printed Electrode and a Pen Electrode assembled from off-the-shelf jumper wire components—and evaluated their performance relative to the discontinued commercial standard.

**Methods:** We conducted in vivo NMES of the left hindlimb ankle dorsiflexors in C57BL/6J mice using three electrode types: a Simple Electrode (previously available as BS4 50–6824, Harvard Apparatus), our novel 3D-Printed Electrode, and our novel Pen Electrode. Contractile torque was recorded during twitch and tetanic contractions, and electrode performance was evaluated based on peak torque values. A repeated measures ANOVA was used to compare torque generated by each animal for the three different types of electrodes.

**Results:** Both custom electrodes generated twitch and tetanic responses comparable to those of the Commercial electrode. No statistically significant differences were observed in peak torque values in either twitch (p = 0.181) or tetany (p = 0.438).

**Conclusion:** Custom-built 3D-printed and Pen-style electrodes offer low-cost, accessible alternatives to discontinued commercial devices for murine NMES studies. These tools can facilitate continued use of electrical stimulation protocols in preclinical neuromuscular research.

## INTRODUCTION

Transcutaneous neuromuscular electrical stimulation (NMES) is a commonly used technique in preclinical studies of muscle function, particularly in rodent models. In mice, NMES protocols are commonly applied to the ankle dorsiflexor muscle group to evaluate contractile function under baseline and post-injury conditions. Central to the delivery of effective stimulation is the use of bipolar electrodes that maintain stable skin contact and deliver current with minimal impedance and artifact.

A previously available bipolar electrode (Simple Electrode, BS4 50–6824 a.k.a., 723742, Harvard Apparatus) was commonly employed for this purpose (Fig. 1A, left) ^1-5^. However, recent correspondence with the manufacturer confirmed that this product is no longer in production. This discontinuation presents a barrier to consistency and reproducibility in ongoing NMES-based studies.

**Figure 1.**
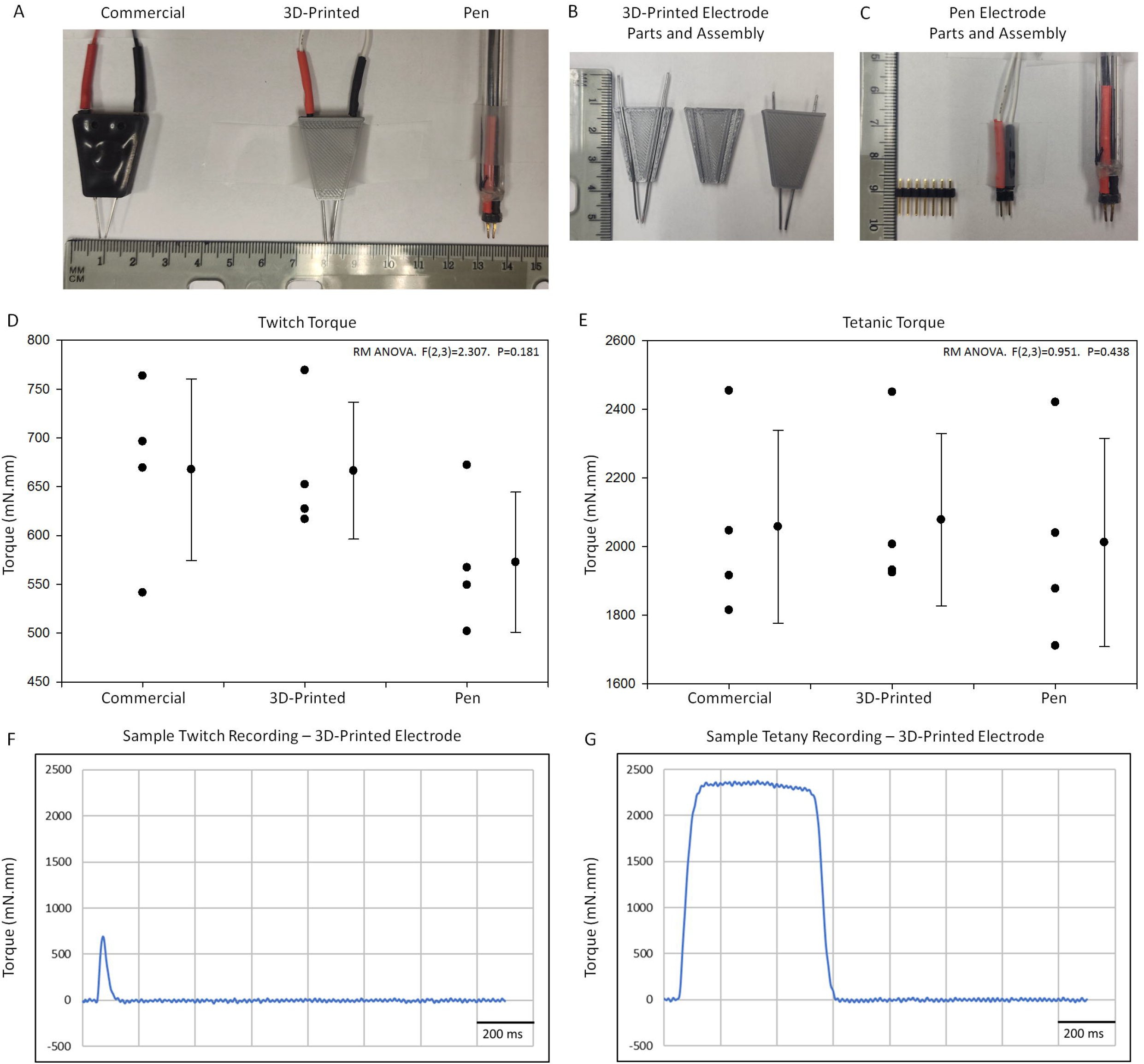
Steps involved in designing and testing custom electrodes for NMES in mice. A. A side-by-side comparison of the appearance of three different electrodes that were tested in this study. B,C. Assembly details of the 3D-Printed and Pen electrodes. D, E. Graphical representation and statistical analysis of twitch and tetanic torque data obtained with the Commercial electrode, 3D-Printed electrode, and Pen electrode during NMES applied to the left hindlimb ankle dorsiflexors of healthy C57BL/6J mice. No statistically significant differences were observed in peak torque values across conditions (twitch p = 0.181; tetany p = 0.438). F,G. Representative twitch and tetanic torque traces obtained with the 3D-Printed electrode.

To address this challenge, we developed and tested two custom-built alternatives (Fig 1A). The first is a 3D-Printed Electrode that incorporates stainless steel wire terminals encased in a rigid, printed housing (Fig. 1A, middle). The second is a Pen Electrode assembled using male jumper wire headers from a commercial electronics kit (Fig. 1A, right). Both designs were developed with accessibility, cost-efficiency, and ease of assembly in mind.

Here, we present a comparative evaluation of these two custom electrodes against the discontinued commercial standard in C57BL/6J mice. We hypothesized that both alternatives would perform equivalently in generating single twitch and fused tetanic torque responses during NMES, thereby offering viable replacements for use in basic and preclinical neuromuscular research.

## METHODS

### Animal Models

All experiments involving live animals were conducted at Wayne State University (Detroit, MI, USA), in accordance with protocols approved by the university’s Institutional Animal Care and Use Committee (IACUC), and in compliance with the Guide for the Care and Use of Laboratory Animals (8th Edition, National Academies Press, 2011) ^6^.

We studied four, male, C57BL/6J mice (Stock No. 000664; The Jackson Laboratory, Bar Harbor, ME, USA), aged 3.5 months. This age and sex were selected to facilitate direct comparisons with previously published datasets ^7^. C57BL/6J mice were chosen as the experimental model due to their well-characterized phenotype and the absence of known neuromuscular or musculoskeletal abnormalities ^7^. All procedures were performed under general anesthesia induced and maintained with inhaled isoflurane (2–5% for induction; 1–4% for maintenance) ^4^. Isoflurane was delivered via nose cone using a tabletop vaporizer and active scavenging system to minimize animal stress and operator exposure^4^. Animals used for this study were not euthanized since they were reserved for additional experiments later.

### Contractile Torque Measurement

We studied the torque produced by the ankle dorsiflexors of the left hindlimb with a custom-built dynamometry rig, as previously described ^1,3^. Mice were exposed to twitch (1 Hz stimulation frequency) and tetanic (125 Hz stimulation frequency) contractions via neuromuscular electrical stimulation (NMES), delivered through each of the three bipolar electrode types evaluated in this study.

For stimulation, we placed the electrode terminals over the lateral aspect of the hindlimb, just distal to the knee joint, targeting the common peroneal (fibular) nerve as it passes around the head of the fibula (video demonstration provided as part of the supplementary materials). Based on our prior work, a subtle bony prominence at this location serves as a reliable tactile landmark for electrode positioning ^4^. We verified precise electrode placement by delivering 1 Hz stimulation and measuring twitch contractions in the tibialis anterior (TA) muscle.

Electrical pulses were generated using an S48 square pulse stimulator connected to a PSIU6 isolation unit (Grass Instruments, West Warwick, RI, USA). NMES parameters were as follows: 0.1 ms pulse duration, 1 pulse train per second, 450 ms train duration. Twitch contractions were elicited with 1 Hz stimulation frequency and fused tetanic contractions were elicited with 125 Hz stimulation frequency (adjusted with the Stimulation Rate dial on stimulator). Custom-written LabView (LabVIEW 2014, National Instruments Corporation, Austin, TX, USA) routines were used to maintain consistent ankle positioning (20° plantarflexion from neutral), trigger the electrical stimulator, and to acquire real-time torque output.

### Electrode Designs

The following three bipolar electrode configurations were evaluated:

#### Commercial Electrode (Standard Control)

This electrode (Simple Electrode, BS4 50–6824 a.k.a., 723742, Harvard Apparatus, Holliston, MA, USA) served as the standard control (Fig. 1A, left). The product has been discontinued, prompting the development and testing of alternative configurations.

#### 3D-Printed Electrode

The custom bipolar electrode (Fig. 1A, middle) was designed using Tinkercad (Autodesk Inc., San Francisco, CA, USA), exported as an STL file, and processed using Ultimaker Cura 5.0 (Ultimaker B.V., Geldermalsen, Netherlands) to generate printer-specific G-code (general 3D printing workflow described earlier ^4^). Each half of the electrode— designed as mirror images—was printed separately using an FLSUN Super Racer (SR) delta 3D printer with PLA filament (HATCHBOX, Rowland Heights, CA, USA). The link to the finalized STL design file is provided as part of the supplementary materials so that users may use, share, and modify the design. The design is covered by a creative commons license, which allows users to use, share and modify the code, but not use it for commercial purposes (CC BY-NC 3.0). The authors respectfully request that this paper be cited as the source.

Each printed half of the electrode housing contained grooves to accommodate 20-gauge stainless steel wires (Cridoz; purchased via Amazon USA)(Fig.1B). Wire segments were manually straightened using plastic-coated pliers and deburred using a metal sanding disc attached to a rotary tool. Following wire placement in the electrode grooves, the two halves of the electrode were aligned and bonded using cyanoacrylate adhesive (Gorilla Super Glue Micro Precise; The Gorilla Glue Company, Cincinnati, OH, USA), forming a rigid, electrically insulated housing (Fig. 1B).

#### Pen Electrode

This electrode (Fig. 1A, right) was assembled using paired male headers from a 9” male-to-female jumper wire kit (Schmartboard Inc., Newark, CA, USA)(Fig.1C). Jumper wires were connected to the headers and threaded through the writing end of a transparent pen barrel (Fig. 1C). The headers were stabilized using hot glue, and the exposed prongs served as the terminals, which contacted the skin during NMES (Fig. 1C).

### Electrode Wiring

To connect each electrode to the isolation unit of the stimulator, we used approximately one meter of 26 AWG 2-conductor red/black wire (Leo Sales Ltd.; available via Micro Center, SKU: 687640). Jumper wire connectors were spliced to the connecting wires to allow for quick and easy electrode changes. The 20-gauge wire terminals of the 3D- Printed Electrode fit snugly into the female connectors of the jumper wires. Standard heat-shrink tubing was used to protect and color-code (red and black) connectors and connections.

### Electrode Testing

We tested each electrode configuration in randomized order on the same animals, with sufficient rest intervals between trials. Multiple twitch and tetanic responses were recorded for each electrode configuration, and the highest recording for each electrode was selected for statistical analysis (Fig 1D, E).

### Data Analysis

We first compiled torque data from twitch and tetanic contractions in Microsoft Excel (Microsoft 365 Apps for Enterprise, Version 2301, Microsoft Corporation, Redmond, WA, USA). We then analyzed maximum twitch and tetanic torque data with SigmaStat 3.5 (Systat Software, San Jose, CA, USA) (Fig. 1D, E). Repeated measures (RM) ANOVA was used to compare conditions (RM, since torque was measured from the same animal’s ankle dorsiflexors across the three different electrodes). We decided *a priori* that p < 0.05 is statistically significant.

## RESULTS

The maximum twitch torque recorded was 668 ± 93, 666 ± 70, 573 ± 72 mN.mm (millinewton-millimeter, units for torque data), respectively, for the Commercial, 3D- Printed, and Pen electrode (Fig. 1D). The group data were statistically indistinguishable [RM ANOVA (F(2,3) = 2.307. P = 0.181], suggesting that all three electrodes are similar in their functional performance with reference to eliciting twitch contractions with 1 Hz stimulation frequency (Fig. 1D). A representative twitch trace elicited with the 3D- Printed electrode is shown in Fig. 1F – the torque traces were qualitatively similar with the Commercial and Pen electrodes – i.e., the twitch torque traces were comparable in their onset, shape, and termination (not shown).

The maximum tetanic torque recorded was 2057 ± 281, 2077 ± 251, 2011 ± 304 mN.mm, respectively, for the Commercial, 3D-Printed, and Pen electrode (Fig. 1E). The group data were statistically indistinguishable [RM ANOVA (F(2,3) = 0.951. P = 0.438], suggesting that all three electrodes are similar in their functional performance with reference to eliciting fused tetanic contractions with 125 Hz stimulation frequency (Fig. 1E). A representative tetanic contraction trace elicited with the 3D-Printed electrode is shown in Fig. 1G – as with twitch, tetanic torque traces were qualitatively similar with the Commercial and Pen electrodes – i.e., the tetanic torque traces were comparable in their onset, shape, and termination (not shown).

## DISCUSSION

This study demonstrates that two easily fabricated custom bipolar electrodes—one 3D- printed and one assembled from off-the-shelf electronics components—can generate torque responses in C57BL/6J mice that are statistically indistinguishable from those elicited by a discontinued commercial electrode.

The loss of access to a reliable commercial electrode presents a significant barrier to reproducibility in mouse NMES protocols. Our alternative designs are not only cost-effective, but also readily assembled using widely available tools and materials. Importantly, both custom electrodes performed reliably in vivo, producing clear and stable twitch and tetany traces with no signal degradation. In 2015, the Commercial electrode cost ∼$40 (USD) – the total cost to produce a 3D-printed or Pen electrode is under $10 (USD), with the jumper wire kit being the major contributor to cost ($8 for 10 pieces, 4 pieces needed for each electrode). For the 3D-Printed electrode, access to a 3D printer and basic operating skills are necessary, but the PLA material is about $25 (USD)/kg and each electrode only uses about 2 g of PLA filament.

While the present study focused on a specific muscle group and a modest sample size, the findings establish proof-of-concept and a pathway for broader application. Future work may explore performance across additional anatomical sites, disease models, and stimulation protocols.

## CONCLUSIONS

In summary, our novel electrode designs offer a practical solution for continued use of NMES paradigms in basic and preclinical neuromuscular research and may enhance accessibility for laboratories that operate on minimal resources.

## Supporting information

3D Printing file information

NMES Demonstration

## ACKNOWLEDGMENTS

This work was supported by NIH R03HD091648, a Pitch Competition Award from the Alliance for Regenerative Rehabilitation Research and Training (AR3T, NIH P2CHD086843), and a subcontract from NIH R01AR079884 (PI: Peter L. Jones) to J.A.R. Additional support was provided by the Wayne State Warrior Funder Program and personal funds from J.A.R. The authors thank the Wayne State University Doctor of Physical Therapy Program for supporting student research and providing training through coursework and research experiences.

