## Supplementary material for "Custom-Built Electrodes Perform Comparably to a Discontinued Commercial Electrode for Neuromuscular Electrical Stimulation in Mice": 3D Printing file information

TINKER CAD LINK:

<https://www.tinkercad.com/things/9EEQGKuZWgi-roche-lab-trapezoidal-bipolar-electrode-v-121224?sharecode=wJ7C56aSKXSdAVgP6K111cjZVsC_w3x1GuVSCGdHaKI>

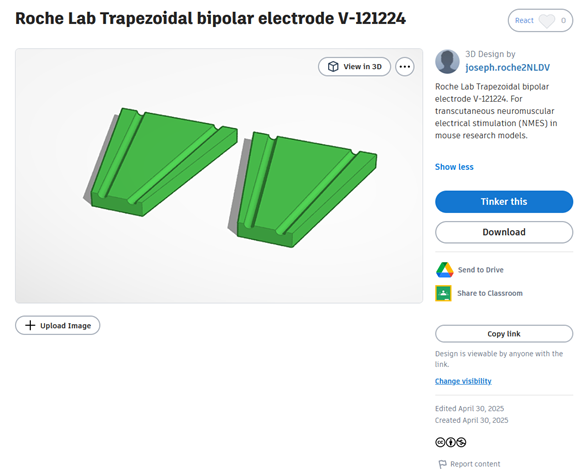

License: <https://creativecommons.org/licenses/by-nc/3.0/>
